# Sexual conflict mediated by ecological sex differences can generate diverse patterns of transgenerational plasticity

**DOI:** 10.1101/846287

**Authors:** Nathan William Burke, Shinichi Nakagawa, Russell Bonduriansky

## Abstract

Transgenerational plasticity (TGP) occurs when the environment experienced by parents induces changes in traits of offspring and/or subsequent generations. Such effects can be adaptive or non-adaptive and are increasingly recognised as key determinants of health, cognition, development and performance across a wide range of taxa, including humans. While the conditions that favour maternal TGP are well understood, rapidly accumulating evidence indicates that TGP can be maternal or paternal, and offspring responses can be sex-specific. However, the evolutionary mechanisms that drive this diversity are unknown. We used an individual-based model to investigate the evolution of TGP when the sexes experience different ecologies. We find that adaptive TGP rarely evolves when alleles at loci that determine offspring responses to environmental information originating from the mother and father are subject to sexually antagonistic selection. By contrast, duplication and sex-limitation of such loci can allow for the evolution of a variety of sex-specific responses, including non-adaptive sex-specific TGP when sexual selection is strong. Sexual conflict could therefore help to explain why adaptive TGP evolves in some species but not others, why sons and daughters respond to parental signals in different ways, and why complex patterns of sex-specific TGP may often be non-adaptive.

## INTRODUCTION

Theory predicts the evolution of anticipatory TGP when environmental conditions experienced by parents and offspring are correlated (Marshall and Uller 2007; Burgess and Marshall 2014). Environmental predictability across generations is thought to allow parents experiencing conditions that fluctuate in space or time to adaptively match the phenotype of their offspring to expected conditions through the transmission of relevant environmental information (Mousseau and Fox 1998; Marshall and Uller 2007; Dey et al. 2016). However, like other forms of plasticity, TGP is not always adaptive. Anticipatory TGP can be costly if the environment of offspring fails to match the parental environment (Crean and Marshall 2009), and some environmental factors appear to induce phenotypic changes that are invariably maladaptive or pathological (Anway et al. 2005; Marshall et al. 2008). Furthermore, mother-offspring conflict can result in induced phenotypes that optimise mothers’ long-term fitness but are suboptimal for individual offspring, and *vice versa* (Uller 2008; Uller and Pen 2011; Kuijper and Johnstone 2018). Although this diversity of effects might explain why evidence for adaptive TGP across studies is generally weak (Uller et al. 2013), existing explanations fail to account for other important patterns of variation in TGP.

An increasing number of studies suggests that TGP is often sex-specific, both in terms of the parent that transfers environmental information and the offspring that responds. Paternal and maternal environments are known to have contrasting effects on offspring (Galloway 2001; Ducatez et al. 2012; Akkerman et al. 2016; Guillaume et al. 2016), and sons and daughters often respond differently to maternal versus paternal information (Pembrey et al. 2006; Dunn et al. 2011; Emborski and Mikheyev 2019; Hellmann et al. 2019), leading to diverse sex-specific patterns of TGP such as mother-daughter, father-son, mother-son and/or father-daughter effects. For example, early-life smoking in human fathers induces larger body mass in sons but not in daughters (Pembrey et al. 2006). In three-spined sticklebacks, predator-exposed mothers produce more cautious offspring regardless of sex, whereas fathers that are similarly exposed produce bolder sons but not bolder daughters (Hellmann et al. 2019). While there is growing interest in the importance of sex-specific TGP in ecology (Uller 2008), conservation biology (Donelson et al. 2018), gerontology (Benayoun et al. 2015), psychology (Bale et al. 2010), neurology (McCarthy et al. 2009), and epidemiology (Skinner et al. 2010), existing models consider only maternal TGP (i.e., environment-induced maternal effects) and typically ignore fathers and sons (e.g., see Uller and Pen 2011; Kuijper and Johnstone 2018). Thus, it remains unclear why maternal and paternal environments often have contrasting phenotypic effects on offspring, and why sons and daughters often respond differently depending on the sex of the parent.

Sex-specific responses to parental environments may reflect underlying differences in the ecology of the sexes. In dioecious organisms, males and females are often selected to utilise environments in different ways as a consequence of sexual dimorphism (Slatkin 1983; Shine 1989; Butler 2007; Butler et al. 2007), which can lead to ecological sex-differences in aggregation (Ruckstuhl and Neuhaus 2006), dispersal (Pusey 1987), foraging (Schoener 1967, 1968; Vitt and Cooper Jr. 1985; Lewis et al. 2002), predation (Schoener and Schoener 1982; Magnhagen 1991), parasitism (Zuk and McKean 1996), nutrient intake (Bearhop et al. 2006; Dalu et al. 2017) and physiological stress (Bale and Epperson 2015). Environmental change can have sex-specific effects as well (Olsson and Van der Jeugd 2002), and such sex-differences are expected to be consistent across generations. Thus, environmental perturbations experienced by mothers (fathers) may predict perturbations experienced by daughters (sons). While correlations between parental and offspring environments are a key condition for the evolution of adaptive TGP (Burgess and Marshall 2014), the implications of sex-specific correlation of environments across generations have not been investigated.

Environmental fluctuations that affect the sexes differently could favour the evolution of sex-specific TGP. If females and males transmit contrasting information to their offspring about the phenotype they should adopt, progeny that pay attention to information from their same-sex parent may gain a fitness benefit because their induced phenotype will match the expected environment for their sex. However, progeny that pay attention to their opposite-sex parent may incur a fitness cost resulting from a mismatch between their phenotype and environment. Assuming that the sexes share a common genetic architecture for response to parental environments, loci that control such responses may therefore be subject to sexually antagonistic selection. Many genetically determined traits that are expressed in both sexes have separate male and female fitness optima (Bonduriansky and Chenoweth 2009). This form of genetic conflict between the sexes (“intralocus sexual conflict”) can result in either an evolutionary stalemate, where each sex expresses the trait sub-optimally (Chippindale et al. 2001), or a resolution, where sex-linked modifiers or duplicated genes (Bonduriansky and Chenoweth 2009; Gallach and Betrán 2011; Vankuren and Long 2018) allow for optimal, sex-specific trait expression (i.e., optimal sexual dimorphism; Ellegren and Parsch 2007). The outcome of conflict at loci responsive to parental information (i.e., stalemate or resolution) could have profound consequences for resulting patterns of TGP. For example, ongoing intralocus conflict might constrain the sexes’ ability to optimise developmental responses, resulting in polymorphic patterns of TGP within species that show little sex specificity. However, resolution of such conflict might generate sexually dimorphic patterns of TGP by releasing the sexes from the constraints of a shared genetic architecture, allowing sons and daughters to attain their sex-specific optima simultaneously.

The potential for sexual conflict mediated by sex-differences in ecology to explain the diversity of observed patterns of TGP remains unknown. Here, using an individual-based model, we investigate the role of sexually antagonistic selection in the evolution of sex-independent and sex-specific TGP under scenarios where loci responsive to environmental information originating from parents are (1) subject to persistent intralocus sexual conflict (‘ongoing conflict’ model); or (2) sex-specific and sex-limited and therefore not subject to sexually antagonistic selection (‘resolved conflict’ model). Each version of the model considers a scenario typical of organisms that show anticipatory TGP in which environmental perturbation—such as higher predation risk (Agrawal et al. 1999), increased temperature (Guillaume et al. 2016), or greater food scarcity (Barrès and Zierath 2016)—causes costly physiological stress. To assess the role of sex-specific ecology in the evolution of TGP, we model scenarios where environmental perturbation affects the sexes in either similar or different ways, such that males, females, both sexes, or neither sex have a higher likelihood of experiencing stress. We assume that individuals cannot alter their own phenotype in response to stress but, instead, transfer information about the changed environment to their offspring via a nongenetic factor. Offspring that carry ‘listening’ alleles can respond to this information from their parents by developing a stress-resistant phenotype that enhances fitness in stressful conditions. Thus, our model adopts an “offspring’s eye view” of TGP, whereby selection acts on sons’ and daughters’ responses (i.e., listening/not listening) to nongenetic information received from the father and mother. (For a similar modelling approach in the context of mother-offspring conflict, see Kuijper and Johnstone 2018). Our model shows how intralocus sexual conflict and its resolution can generate the diversity of patterns of TGP seen in nature, including adaptive, non-adaptive, sex-specific and sex-independent effects, thereby providing a new, unifying framework for the evolution of TGP.

## THE MODEL

### Life cycle and stress ecology

Our individual-based model considers a dioecious, bi-sexual population with discrete generations distributed over 21 × 21 patches in a torus-shaped world, with no limitation on the number of individuals per patch. Patches vary in the amount of food they hold, which is determined at the start of each generation by assigning a randomly sampled integer to each patch from the discrete uniform distribution U{0, 2}. Food resources are fixed for the duration of each generation and cannot be reduced or depleted. Each generation begins with the simultaneous emergence of adult individuals with 20-timestep lifespans, distributed randomly across patches. In every timestep, adults move randomly to one of the eight patches bordering their current patch and consume food resources, with the amount consumed dependent on the amount of resources available in that patch. The cumulative amount of resources consumed over the lifetime contributes to condition at mating/reproduction. Individuals also experience the density of conspecifics in each patch visited, and this density contributes to costly stress (with individual susceptibility to stress determined by inherited nongenetic factors, sex, and ‘listening’ genotype, as described below). For simplicity, we assume that adults do not alter their behaviour in response to food or conspecific density. In the final time-step, adults mate, reproduce, and die. Reproductive success is affected by condition at the end of the lifetime, which is an increasing function of the amount of resources accumulated and a decreasing function of the extent of phenotypic mismatch to stressful conditions, as described below. Figure 1 provides a visual summary of the lifecycle.

**FIGURE 1.**
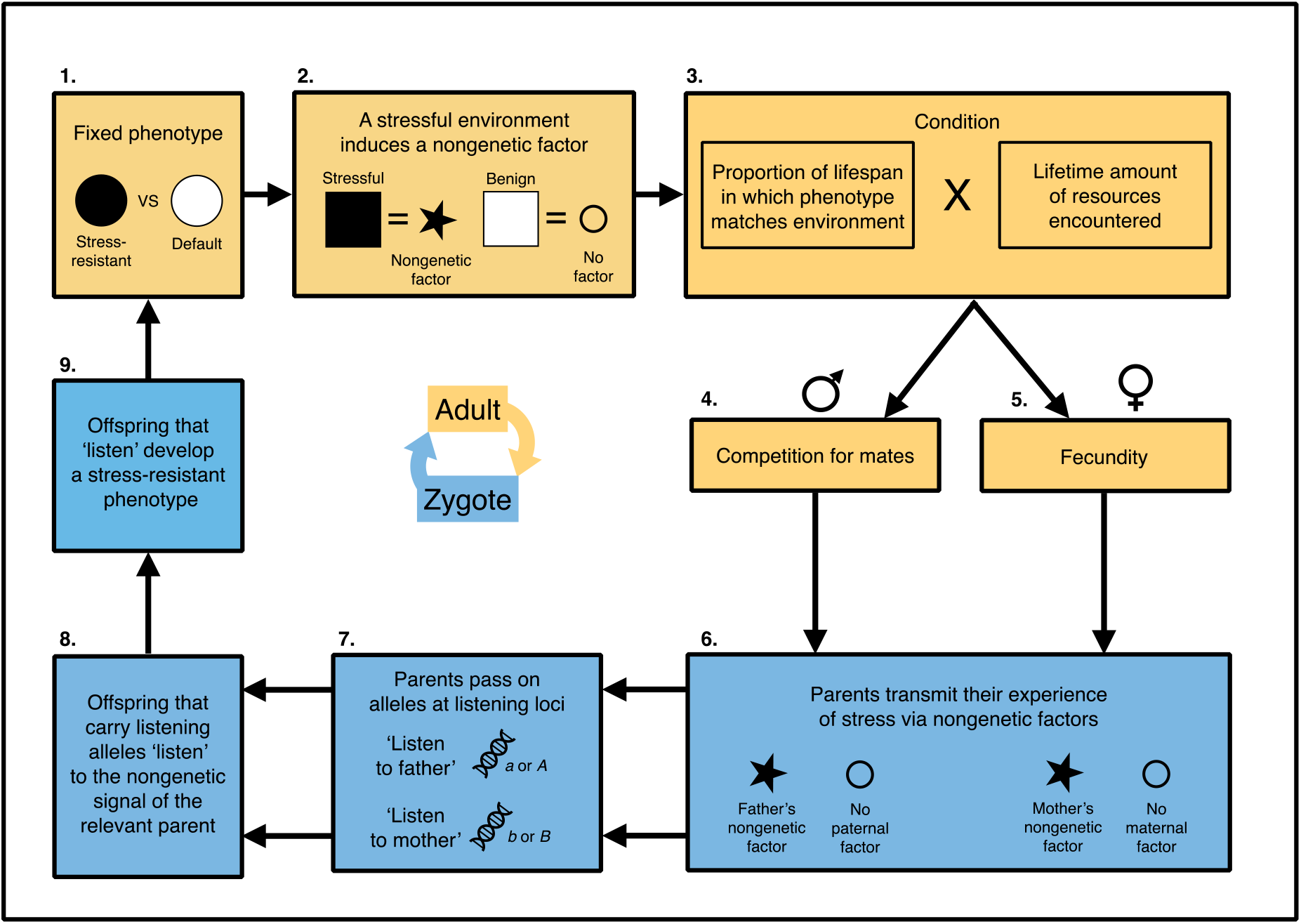
Lifecycle of organisms in the ongoing conflict model. At the start of the lifecycle, adult individuals emerge with a default phenotype suited to benign conditions that is fixed for the lifetime (1). A stress-resistant phenotype is also possible but must be induced transgenerationally. Individuals experience stress when they interact with too many conspecifics. Stress induces the production of a nongenetic factor (2), which is later transferred to offspring. The extent to which an individual’s phenotype matches its experience of stress determines its condition (3). For males, condition determines mating success (4), whereas for females, condition determines fecundity (5). Females mate with the most competitive male in their mating neighbourhood, or with a random male as a control. At reproduction, both parents faithfully pass on their nongenetic factors (6) as well as listening alleles (7). Offspring that carry listening alleles utilise nongenetic information from parents (8) to develop a stress-resistant phenotype (9). Offspring without listening alleles ignore parental information and develop a default phenotype suited to benign conditions (1). The same lifecycle applies to the resolved conflict model except that sex-specific listening loci *C*, *D*, *E* and *F* replace loci *A* and *B*.

Our model assumes that adults experience stress when interacting with conspecifics (Champagne 2010; Gudsnuk and Champagne 2012; Dantzer et al. 2013), and that males and females encounter different levels of stress due to one of the sexes being more likely than the other to interact with surrounding individuals in the patch. That is, we assume that the density of conspecifics encountered by individuals within a patch is random, but that the rate of interaction with the individuals encountered is sex-specific. Consistent with the physiology of stress tolerances in animal systems (Pörtner 2002), we assume that stress (a binary state) is triggered when the cumulative number of intra-patch conspecific encounters that an individual has experienced in its life so far (*K*_*ind*_) surpasses a threshold (*α*); that is, when *K*_*ind*_ ≥ *α*_*m*_ (or *K*_*ind*_ ≥ *α*_*f*_), where *α*_*m*_ (*α*_*f*_) is the number of encounters that males (females) can withstand without becoming stressed. Thus, once triggered, stress persists for the rest of the lifetime. Thresholds *α*_*m*_ and *α*_*f*_ are allowed to vary for a fixed high or low value (80 and 8, respectively), such that the sexes can have a similar or different probability of experiencing stress. Thus, a high *α*_*m*_ and low *α*_*f*_ threshold would represent a scenario where males rarely interact with surrounding conspecifics (and therefore infrequently experience costly stress), while females regularly interact with conspecifics (and therefore often experience costly stress). Such sex-differences in the probability of conspecific interaction are likely to arise in natural systems as a consequence of innate sex-differences in life-history strategies, such as females seeking high quality food resources, or males searching for mates, aggregating in leks, or avoiding competitors (Clutton-Brock 1989). While we focus on TGP triggered by high rates of interaction with conspecifics, our model generalizes to any type of adaptive TGP that evolves in response to environmental challenges that may be sex-specific, such as predation, parasitism, thermal stress, nutrient intake, or starvation.

Models of anticipatory TGP often assume periodic fluctuations between environmental states (Uller and Pen 2011; Kuijper and Johnstone 2018). For simplicity, we consider a single period of sex-specific stress. However, simulations with multiple periodic shifts in stress produce very similar results (see Supplementary Figure S1).

### Transgenerational plasticity

Given the importance of constraints on phenotypic plasticity for the evolution of TGP (Kuijper and Hoyle 2015), we assume that adults cannot plastically change their own phenotype to cope with stress but transmit to offspring a stress-induced, nongenetic factor that has the potential to alter offspring development. We make no assumptions about the type of factor involved or whether stressed mothers and fathers necessarily transmit the same factor. Our model simply assumes that changes in nongenetic factors—whether transcription factors (Tao et al. 2017), small RNAs (Chen et al. 2016), DNA methylation (Weaver et al. 2004; Herman and Sultan 2016), histone modification (Morgan et al. 2005), nutrients (Blount *et al.* 2000), hormones (Groothuis *et al.* 2005), antibodies (Hasselquist & Nilsson 2009), parental care (Eggert et al. 1998), or something else—are consistently triggered by stress, that offspring faithfully inherit these factors from stressed parents, and that such factors carry information about the stressfulness of the parental environment. Offspring that inherit stress-induced factors and also carry appropriate ‘listening’ alleles develop into individuals with a stress-adapted phenotype (see Genetic architecture, below). We assume that offspring are capable of responding differently to factors received from the mother versus the father, as suggested by reported examples of sex-specific TGP (Pembrey et al. 2006; Zizzari et al. 2016; Hellmann et al. 2019), and that the phenotypic effect of receiving factors from both parents is the same as receiving a factor from one (i.e., no additive or multiplicative effects), such that offspring that receive conflicting information listen to the signal of the stressed parent. However, offspring that do not carry listening alleles are unable to alter their development in response to nongenetic information, and instead develop the default, non-resistant phenotype.

Since mismatch between phenotype and environment is often modelled as a fitness cost of TGP (e.g., Kuijper and Johnstone 2018), we include this cost in our model but allow it to be sex-specific. Individuals with the default phenotype are considered well-matched if they experience benign conditions (i.e., *K*_*ind*_ < *α*_*m*_ or *K*_*ind*_ < *α*_*f*_) and mismatched if they experience stressful conditions (i.e., *K*_*ind*_ ≥ *α*_*m*_ or *K*_*ind*_ ≥ *α*_*f*_); conversely, individuals with the stress-resistant phenotype are considered well-matched if they experience stressful conditions (i.e., *K*_*ind*_ ≥ *α*_*m*_ or *K*_*ind*_ ≥ *α*_*f*_) and mismatched if they experience benign conditions (i.e., *K*_*ind*_ < *α*_*m*_ or *K*_*ind*_ < *α*_*f*_). Thus, the level of match (*∂*_*ind*_) to stressful conditions is phenotype-dependent, such that *∂*_*ind*_ = 1 − *δ*_*ind*_ for default phenotypes and *∂*_*ind*_ = *δ*_*ind*_ for stress-resistant phenotypes, where *∂*_*ind*_ is the proportion of the lifetime spent well-matched, and *δ*_*ind*_ is the proportion of the lifetime spent in a stressed state. Because phenotype-environment mismatch can be maladaptive (Kettlewell 1956; Zimova et al. 2016), we assume that phenotypic match affects an individual’s ability to accumulate resources, and therefore its condition at mating/reproduction (*C*_*ind*_), such that *C*_*ind*_ = *∂*_*ind*_. *φ*_*ind*_, where *φ*_*ind*_ is the total quantity of resources encountered during the lifetime. Thus, the greater the proportion of time spent matched, the greater the amount of resources an individual is able to accumulate, and the higher its condition at mating/reproduction. Condition determines female fecundity (*W*_*ind*_) and male competitiveness (*Q*_*ind*_), such that *W*_*ind*_ = *C*_*ind*_ and *Q*_*ind*_ = *C*_*ind*_. These positive linear relationships reflect the strong condition dependence of male secondary sexual traits (Andersson 1982) and female fecundity (Honěk 1993) in natural systems.

### Selection, mating and reproduction

Selection is sex-specific, with sexual selection on males occurring during the mating phase, and fecundity selection on females occurring during the reproduction phase. In both cases, phenotypic match (*∂*_*ind*_), which determines *Q*_*ind*_ and *W*_*ind*_, is the phenotypic trait directly subject to selection.

At the end of each generation, females mate once with the male in their mating neighbourhood that has the highest condition (i.e., the ‘best male’), or a random male as a control (‘random mating’). Thus, selection acts on both sexes under best male settings, but only on females when mating is random. The explicit spatial structure of our model allows us to investigate the consequences of sexual selection intensity by varying the size of the mating neighbourhood (1 patch, 9 patches, 25 patches, 49 patches), with larger neighbourhoods generating a larger skew in male mating success (i.e., more intense sexual selection).

Reproduction occurs immediately following mating, within the same timestep. For simplicity, we assume no male contribution to fecundity. Females produce a total of *W*_*ind*_ = *C*_*ind*_ offspring (to the nearest integer), with sons and daughters equally likely to be produced. The total number of offspring, *n*_*gen*_, produced at the end of a generation is always more than can survive (i.e., greater than the global carrying capacity, *P*). To maintain a stable population size, *n*_*gen*_ − *P* offspring are killed randomly before eggs hatch. We assume *P* = 1000 for all simulations. The lifecycle repeats following the death of all parents and the simultaneous emergence of offspring.

### Genetic architecture

Although the genetic architecture of offspring response to nongenetic factors is poorly understood, our assumption of genetic control of this response is motivated by empirical findings that TGP can depend on offspring genotype (Cayuela et al. 2019). Our ongoing conflict model assumes two listening loci that determine offspring responses to information from stress-exposed fathers (locus *A*) and mothers (locus *B*). Phenotypic responses of offspring are necessarily sex-independent in this version of the model because males and females share the same genetic architecture for listening. Two alleles segregate at each locus (wildtype: *a* and *b*; mutants: *A* and *B*), with additive effects on the probability of listening (0% chance of listening: *aa* and *bb*; 50% chance of listening: *Aa* and *Bb*; 100% chance of listening: *AA* and *BB*) (see Table 1 for a summary of phenotypic effects). Note that listening alleles determine whether offspring respond to nongenetic factors, not whether parents transmit those factors.

**TABLE 1.** Phenotypic effects of listening genotypes. When sexual conflict is ongoing, listening alleles have sex-independent effects on offspring responses to maternal and paternal information; whereas when sexual conflict is resolved, listening alleles have sex-specific effects on offspring responses. Note that an individual’s overall genotype is determined by the alleles it carries at each locus.

|  | Locus | Genotype | Phenotype |
| --- | --- | --- | --- |
| Ongoing conflict | *A* | *aa* | Offspring do not listen to father's information |
|  |  | *Aa* | Offspring listen to father's information 50% of the time |
|  |  | *AA* | Offspring listen to father's information 100% of the time |
|  | *B* | *bb* | Offspring do not listen to mother's information |
|  |  | *Bb* | Offspring listen to mother's information 50% of the time |
|  |  | *BB* | Offspring listen to mother's information 100% of the time |
| Resolved conflict | *C* | *cc* | Sons do not listen to father's information |
|  |  | *Cc* | Sons listen to father's information 50% of the time |
|  |  | *CC* | Sons listen to father's information 100% of the time |
|  | *D* | *dd* | Sons do not listen to mother's information |
|  |  | *Dd* | Sons listen to mother's information 50% of the time |
|  |  | *DD* | Sons listen to mother's information 100% of the time |
|  | *E* | *ee* | Daughters do not listen to father's information |
|  |  | *Ee* | Daughters listen to father's information 50% of the time |
|  |  | *EE* | Daughters listen to father's information 100% of the time |
|  | *F* | *ff* | Daughters do not listen to mother's information |
|  |  | *Ff* | Daughters listen to mother's information 50% of the time |
|  |  | *FF* | Daughters listen to mother's information 100% of the time |

The resolved conflict model assumes that listening loci are duplicated and sex-limited: locus *C* controls sons’ listening to fathers, locus *D* controls sons’ listening to mothers, locus *E* controls daughters’ listening to fathers, and locus *F* controls daughters’ listening to mothers. The loci are unlinked, and two alleles segregate at each locus (wildtype: *c*, *d*, *e*, and *f*; mutant: *C*, *D*, *E*, and *F*), with additive effects on the probability of listening (0% chance of listening: *cc*, *dd*, *ee* and *ff*; 50% chance of listening: *Cc*, *Dd*, *Ee* and *Ff*; 100% chance of listening: *CC*, *DD*, *EE* and *FF*) (see Table 1 for a summary of phenotypic effects).

### Simulations

In all simulations, mutations at listening loci are introduced haphazardly at the start of generation 25 following a short burn-in period to allow population dynamics to stabilise, such that wildtype alleles are replaced by listening alleles, and *vice versa*, at an ongoing per-locus per-timestep rate *r* (fixed at *r* = 0.001).

To assess the role of sexual conflict in the evolution of sex-independent and sex-specific listening, we varied parameters controlling male and female sensitivity to conspecific encounter, *α*_*m*_ and *α*_*f*_, and the intensity of sexual selection on males, for both the ongoing conflict and resolved conflict models. We ran 40 simulations per parameter combination for 1000 generations and recorded allele frequencies at each listening locus at the end of each run. We considered listening to have evolved if the frequency of a listening allele was ≥ 0.95 at the end of a simulation run, and polymorphisms in listening to have occurred if the frequency of a listening allele remained stable and consistently greater than 0 and less than 0.95 for the duration of simulations. To understand how selection was acting on the sexes, we also recorded the sex-specific distribution of phenotypic matching for each parameter combination at the end of each run.

## RESULTS

### Ongoing conflict

The introduction of listening alleles at loci that control embryos’ ability to respond to the epigenetic signal from their father (locus *A*) and mother (locus *B*) leads to four evolutionary outcomes: paternal TGP (allele *A* fixes), maternal TGP (allele *B* fixes), both types of TGP (alleles *A* and *B* fix), or no TGP (neither *A* nor *B* fixes). The relative level of stress experienced by each sex largely determines the probability of these outcomes (see Figure 2a). When males experience higher stress than females, listening to fathers (allele *A*) is favoured in sons but disfavoured in daughters because the stress-induced phenotype that listening permits improves the phenotypic match (and, hence, fitness) of sons but decreases the match of daughters. Listening to mothers (allele *B*) is neutral in this scenario, since mothers rarely experience stress. However, despite the evolutionary benefit to males of listening to fathers, sexual conflict at locus *A* largely inhibits the fixation of the *A* allele, which codes for paternal TGP, because the benefit of paternal TGP to sons is not enough to overcome the cost of paternal TGP to daughters (hence, the paucity of outcomes where the *A* allele approaches fixation in Figure 2a). A similar but converse situation occurs when females experience more stress than males (Figure 2a). Thus, despite the benefit that listening provides to the stressed sex, sexual conflict at listening loci constrains the evolution of TGP because of the costs that listening imposes on the opposite sex.

**FIGURE 2.**
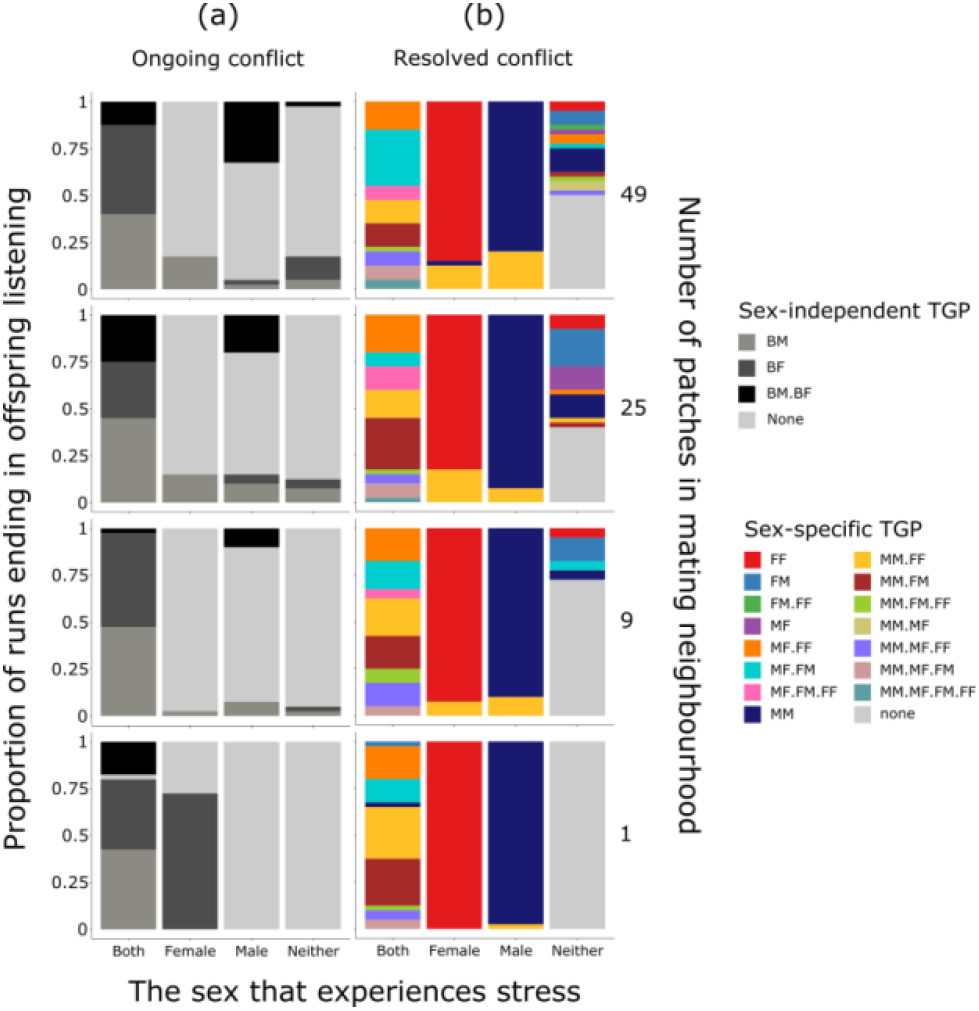
Sex-independent listening (a) and sex-specific listening (b) Graphs on the left (a) show the proportion of runs of the best male version of the ongoing conflict model ending in sex-independent paternal TGP (dark grey), maternal TGP (medium grey), both types of TGP (black), and no TGP (light grey) at generation 1000. Graphs on the right (b) show the proportion of simulation runs of the best male version of the resolved conflict model ending in sex-specific TGP at generation 1000. Note that sexual selection occurs under best male settings and increases in intensity with number of patches in the mating neighbourhood. Colours in the legend represent all possible combinations of listening. The first letter in each double-letter code represents the sex of the offspring that does the listening (M = males (i.e., sons); F = females (i.e., daughters); B = both sexes), and the second letter represents the sex of the parent that is listened to (M = males (i.e., fathers); F = females (i.e., mothers)). Thus, for example, BM indicates a sex-independent paternal effect (i.e., sons and daughters both listening to fathers), and MF indicates a son-specific maternal effect (i.e., sons listening to mothers). Colours with more than one double-letter code indicate runs where more than one combination of sex-independent or sex-specific TGP evolves. Runs were counted as ending in TGP if listening allele frequencies were ≥ 0.95. In (a), when one parent experiences higher stress than the other, intralocus conflict is generated between sons and daughters over whether to pay attention to information received from the stressed or unstressed parent. In (b), sex-specific listening loci allow sexual conflict to be resolved via the evolution of sex-specific TGP.

The sexes’ evolutionary interests in listening diverge when one sex experiences higher stress than the other and, in our model, this conflict manifests as sex-differences in *∂*_*ind*_ (see Figure 3a). Sexual selection on males has a strong influence on the distribution of *∂*_*ind*_ and hence patterns of listening under sexual conflict. Indeed, the relative strength of selection on the sexes modulates the effect of intralocus conflict on listening outcomes. In the absence of sexual selection, fecundity selection maximises female matching regardless of the parent that experiences stress (see Supplementary Figure S2), which leads to optimal listening outcomes for females (i.e., the evolution of maternal TPG when females experience higher stress, and no TGP when males experience higher stress; see Supplementary Figure S3). However, sexual selection on males counteracts this effect. When sexual selection and fecundity selection are similar in strength but act in opposite ways on stress-resistance, stable sex-specific listening polymorphisms evolve (see Figure 4). These intermediate allele frequencies are associated with bimodal sex-differences in *∂*_*ind*_ whereby each sex exhibits both high and low matching (see Figure 3a), reflecting an evolutionary stalemate between the sexes—a predicted outcome of intralocus sexual conflict (Bonduriansky and Chenoweth 2009). Increasing the intensity of sexual selection gives males a greater edge in the conflict, as evidenced by the reduced number of simulations ending in maternal TGP when females experience greater stress (Figure 2a).

**FIGURE 3.**
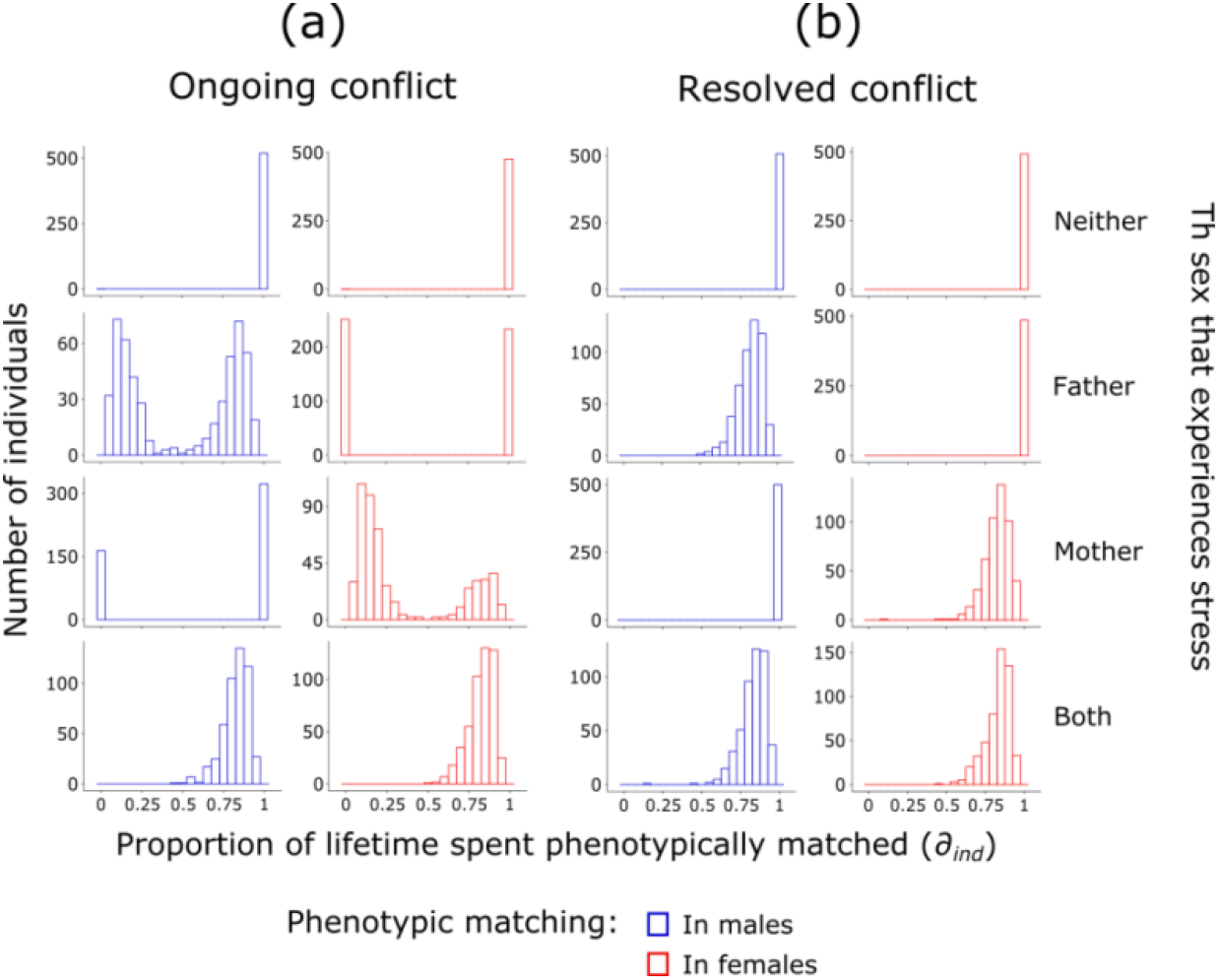
Patterns of phenotypic matching (*∂*_*ind*_) Histograms show the outcome of selection on males (blue) and females (red) at the end of a single simulation run of the best male version of the ongoing conflict model (a) and resolved conflict model (b) at generation 1000. Note that sexual selection occurs under best male settings. In (a), sex-differences in the distribution of matching is indicative of intralocus sexual conflict and occurs when one parent experiences higher stress than the other. These divergent distributions become bimodal due to the opposing action of sexual selection on males and fecundity selection on females. The resolution of sexual conflict in (b) allows each sex to achieve maximal matching at the same time. For the unstressed sex in both (a) and (b), distributions of *∂*_*ind*_ show little variation around the peak because individuals of this sex rarely encounter more conspecifics than their sex’s encounter threshold (*α*_*m*_ or *α*_*f*_), and so are almost always uniformly well-matched. Other settings: mating neighbourhood size = 25 patches.

**FIGURE 4.**
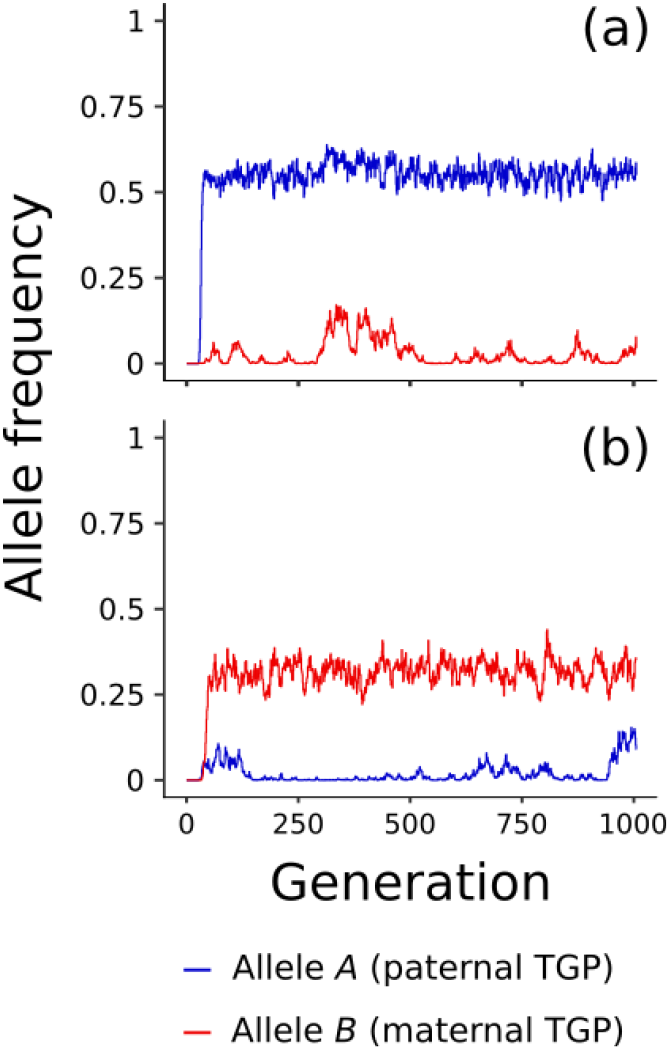
Opposing selection on the sexes generates listening polymorphisms. Graphs show the frequencies of allele *A* (paternal TGP; blue) and allele *B* (maternal TGP; red) for a single simulation run of the best male version of the ongoing conflict model when males experience higher stress (a) and females experience higher stress (b). Note that sexual selection occurs under best male settings. Listening polymorphisms are consistently maintained for hundreds of generations. Other settings: mating neighbourhood size = 25 patches.

In simulations in which neither parent is stressed, listening alleles are neutral and there is no selection for listening (Figure 2a). However, in cases where mothers and fathers are both stressed, listening to either parent is favoured in both sons and daughters, and the particular form of TGP that evolves (paternal, maternal or both) is determined by whichever alleles happen to spread faster by chance (Figure 2a).

### Resolved conflict

When intralocus sexual conflict is resolved through the duplication and sex-limitation of listening loci (Gallach and Betrán 2011), the sexes can pursue optimal listening strategies free from the constraints of a shared genetic architecture. This resolution allows for simulations of the resolved conflict model to end in any combination of sex-specific maternal and paternal TGP.

The resolution of intralocus conflict is characterised by higher values of *∂*_*ind*_ (Figure 3b) and optimal sex-specific listening outcomes (Figure 2b) for both sexes. When females experience higher stress than males, fecundity selection drives daughters to listen exclusively to mothers (Figure 2b). Conversely, when males experience higher stress than females, sexual selection drives sons to listen exclusively to fathers (Figure 2b). These patterns evolve because sexual selection and fecundity selection are able to act independently on son-specific and daughter-specific listening alleles, respectively. The resolution of sexual conflict allows for the adaptive evolution of these simple mother-daughter and father-son effects in a similar way to parent-of-origin effects on gene expression (Day and Bonduriansky 2004).

However, our model also shows how more complex patterns of TGP involving multiple parent-offspring effects can evolve. Diverse sex-specific listening outcomes can evolve when stress is experienced by (1) both parents (representing adaptive effects) or (2) neither parent (representing non-adaptive effects) (see Figure 2b). In the first case (1), diverse combinations result from the random fixation of equally beneficial alleles: if both parents send similar stress signals, sons and daughters can reap similar benefits by listening to either parent. Very few simulations in this parameter space end in only one sex listening because sexual selection and fecundity selection simultaneously favour listening in both males and females. In the second case (2), stochastic patterns of sex-specific listening evolve via random fixation of neutral alleles driven by sexual selection. As expected, no TGP evolves if both parents are unstressed and sexual selection is weak (Figure 2b). However, intensifying sexual selection leads to increased diversity of sex-specific combinations of TGP. Indeed, 11 out of the 15 possible listening combinations evolve at the highest sexual selection intensity (Figure 2b). This diversity occurs because the skew in male mating success generated by intense sexual selection substantially reduces effective population size, causing neutral alleles carried by the most successful males to reach fixation via an effect analogous to hitchhiking.

## DISCUSSION

We modelled the evolution of environmentally induced TGP for stress-resistance within a heterogeneous environment where mothers and fathers can experience different levels of stress that are correlated within sex across generations. We then asked how sex-differences in stress ecology might drive selection on alleles that allow offspring to respond (‘listen’) to parental information and develop a stress-resistant phenotype when the genetic architecture of listening is either shared between the sexes or independent of sex. Our model allowed us to investigate the extent to which intralocus sexual conflict and its resolution could explain the diverse patterns of sex-specific and sex-independent TGP in natural systems. Our model also enabled us to ask how selection acting more strongly on one sex than the other affects listening outcomes.

Our simulations produced five key findings. First, if both sexes share the same genetic architecture for listening and one sex experiences a greater probability of stress than the other, intralocus sexual conflict will inhibit the spread of listening alleles. The resulting allele frequencies for each sex will then be determined by the relative strength of selection on males and females: stronger selection on one sex will skew listening outcomes more toward that sex’s optimum, whereas selection of similar strength will generate bimodal fitnesses and stable listening polymorphisms for each sex. Second, sex-independent listening outcomes, where offspring listen to parents regardless of their own sex, will occur most frequently when sons and daughters gain equal benefit from listening (i.e., when both sexes experience stress and therefore have the same evolutionary interest in listening to stress signals). Third, the resolution of intralocus sexual conflict through duplication and sex-limitation of listening loci, which allows each sex to optimise listening independent of the other sex, will generate daughter-specific maternal TGP when females experience greater stress and son-specific paternal TGP when males experience greater stress. Fourth, complex patterns of sex-specific listening, whereby sons and daughters independently listen to one or both parents, can evolve if conflict is resolved and stress is high for both sexes. Under these settings, all listening alleles have equal benefit to either sex, and the allele that spreads most quickly by chance will go to fixation. Fifth, complex listening combinations can also evolve if conflict is resolved and neither sex experiences stress. In this situation, all listening alleles are neutral and strong sexual selection can drive random combinations of listening alleles to fixation via an effect analogous to genetic hitchhiking.

Our results suggest that a range of phenomena reported in the empirical TGP literature could be explained by sexual conflict. Maternal and paternal environments are known to affect the phenotypes of sons and daughters in different ways (Pembrey et al. 2006; Von Engelhardt et al. 2006; Dunn et al. 2011; Kruuk et al. 2015; Emborski and Mikheyev 2019; Hellmann et al. 2019), and our model provides a potential explanation for this intriguing pattern of TGP. The key assumption of our model that generates sex-biased outcomes is the sex-specificity of environmental stress. Existing models of TGP also assume environmental correlations spanning multiple generations (Motro 1983; Godfray and Parker 1991; Kilner and Hinde 2008; Uller and Pen 2011; Kuijper and Johnstone 2018), but such models overlook the contribution of the paternal environment to offspring phenotypes (i.e., paternal effects; Rando 2012; Crean and Bonduriansky 2014; Soubry et al. 2014), as well as the sex-specificity of offspring responses to parental information (e.g., Pembrey et al. 2006; Zizzari et al. 2016; Hellmann et al. 2019). By incorporating both sexes and sex-specific environments, our model shows how differences in parental signals driven by sex-differences in ecology can lead to the evolution of sex-biased TGP.

Our simulations also suggest that sex-specific patterns of TGP may often be non-adaptive. One of the surprising results from our model was the diversity of sex-specific outcomes that evolved when sex-limited listening alleles were selectively neutral. Sexual selection on males facilitated the spread of neutral listening alleles via a kind of hitchhiking effect, resulting in diverse combinations of sex-specific TGP. This result suggests that organisms characterised by intense sexual selection may be more likely to exhibit complex patterns of sex-specific TGP. Other processes that promote genetic drift, such as genetic bottlenecks, founder effects and genetic hitchhiking, could generate similar results. The diversity of sex-specific outcomes that evolved in our model suggests that much of the sex-specific variation in TGP in natural systems (e.g., Pembrey et al. 2006; Dunn et al. 2011; Emborski and Mikheyev 2019) may be non-adaptive. A finding of diverse and unpredictable sex-specific TGP following experimental manipulation of the intensity of sexual selection would lend empirical support to this non-adaptive explanation.

Our model may additionally help to explain why many studies fail to observe adaptive TGP despite *a priori* predictions of an adaptive benefit (see reviews by Uller et al. 2013; Heard and Martienssen 2014). Existing models suggest that mother-offspring conflict—where the fitness optima of mothers and offspring with respect to offspring phenotype are different and cannot be achieved simultaneously (Trivers 1974)—can lead to the breakdown of TGP under certain conditions (Uller and Pen 2011; Kuijper and Johnstone 2018). Our simulations show that conflict between the sexes—which is ubiquitous in bisexual taxa (Arnqvist and Rowe 2005)—could also inhibit the fixation of alleles for TGP. Intralocus sexual conflict could therefore be a powerful and taxonomically widespread explanation for the absence of adaptive TGP in many taxa, just as intralocus sexual conflict can constrain the evolution of sexual dimorphism in many other traits (Bonduriansky and Chenoweth 2009). Indeed, the ubiquity of sex-specific ecologies in natural systems (Schoener 1967, 1968; Schoener and Schoener 1982; Vitt and Cooper Jr. 1985; Pusey 1987; Shine 1989, 1991; Magnhagen 1991; Zuk and McKean 1996; Temeles et al. 2000; Mysterud 2000; Lewis et al. 2002; Olsson and Van der Jeugd 2002; Bearhop et al. 2006; Ruckstuhl and Neuhaus 2006; Butler 2007; Butler et al. 2007; Wearmouth and Sims 2008; Bale and Epperson 2015; Dalu et al. 2017; Fryxell et al. 2019), and the conflicting signals that parents in such ecologies are likely to send to offspring, could prevent the evolution of adaptive TGP, unless the intersexual genetic correlation for listening can be reduced through changes in genetic architecture involving locus duplication or sex-linked modifiers that enable sex-specific listening. Our model also shows that intralocus sexual conflict could maintain genetic variation for listening via the evolution of stable listening polymorphisms. This is a predicted outcome of non-random mating with respect to traits subject to sexual conflict (Härdling and Bergsten 2006), which explains why polymorphisms evolve in our simulations only when sexual selection is present.

Empirical studies indicate that TGP can be induced by a variety of environment factors, including temperature (Salinas and Munch 2012), light (Galloway and Etterson 2007), food availability (Vijendravarma et al. 2010), food quality (Bonduriansky and Head 2007), water availability (Herman and Sultan 2016), predation risk (Sheriff et al. 2010), pathogen exposure (Sadd et al. 2005) and parasite load (Lefèvre et al. 2010). Although we specifically model environmental perturbation as stressful, this assumption does not reduce the generality of our conclusions as the sexes may experience benign or resource-rich environments differently as well. For example, offspring traits such as body size that can be enhanced by high parental condition (Qvarnström and Price 2001; Crean and Bonduriansky 2014) often have sex-specific optima (Badyaev and Martin 2000), and therefore alleles for listening to such cues may also be subject to sexual conflict. Our findings also highlight the importance of accounting for sex-specific variation in ecology when motivating TGP experiments and interpreting results, as failure to do so could lead to potentially misleading conclusions (see also Kruuk et al. 2015).

Our model could be tested empirically by assessing the expected match between fitness outcomes of phenotypic responses for each sex and the type of TGP that is observed. For example, an absence of a clear signal of TGP associated with contrasting fitnesses between the sexes with respect to offspring phenotype would suggest that fixation of alleles at loci responsive to parental environments is inhibited by ongoing intralocus conflict. Further support for such a conclusion would be found if experimentally decreasing the intensity of selection on one sex results in the evolution of patterns of TGP that are more aligned to the fitness interests of the opposite sex. Assessment of the genetic architecture of TGP in natural systems could also prove valuable. There is evidence that TGP has a genetic basis (Stjernman and Little 2011; Herman and Sultan 2016; Cayuela et al. 2019), but the extent to which the expression of TGP depends specifically on genes that regulate sex-specific development remains an open question. Genetic analyses could determine whether sex-specific TGP, where sons (daughters) respond exclusively to paternal (maternal) environments (e.g., Pembrey et al. 2006; Zizzari et al. 2016; Hellmann et al. 2019), is mediated by listening alleles that are sex-limited in offspring.

The strong inhibiting effect of intralocus conflict on offspring listening that we observed in our model suggests that TGP may be especially widespread in organisms for which sexual conflict is absent or weak. For example, TGP may be particularly common in asexual lineages in which adaptive maternal effects can evolve in the absence of males. However, mother-offspring conflict may be a more prevalent factor in such lineages (Kuijper and Johnstone 2018). Similarly, monogamous species may have a greater capacity for TGP if sexual conflict is relatively weak in such taxa (Holland and Rice 1999; Martin and Hosken 2003) and if monogamy is associated with reduced sex-specificity of ecology (Fryxell et al. 2019; Giery and Layman 2019). Comparative analyses of the incidence of TGP in sexual versus asexual taxa, and monogamous versus polyandrous taxa, could shed light on these questions.

Future work could usefully extend our model by investigating how the evolution of alternative strategies for coping with environmental stress in parents, such as phenotypic plasticity in development (Denver 1997), morphology (Gratani 2014) or behaviour (Badyaev 2005), or offspring sex-ratio adjustment (Nager et al. 1999; Rosenfeld and Roberts 2004), modulates or inhibits the evolution of TGP. Assessment of the effect of temporal heterogeneity in sex-differences, and error in nongenetic signals and offspring responses, may also be fruitful.

Our model suggests a number of novel predictions. First, species in which the sexes experience substantially different ecologies may be more likely to exhibit non-adaptive or polymorphic TGP due to ongoing intralocus conflict at listening loci, or, alternatively, adaptive sex-specific TGP if conflict is resolved through the evolution of sex-specific listening loci, with mother-daughter (father-son) effects most likely when females (males) experience greater stress. Second, sex-independent TGP may be common in species in which males and females experience similar ecologies and no sexual conflict over listening. Third, complex patterns of sex-specific TGP are likely to be non-adaptive, and may be most common in species in which males are subject to strong sexual selection. Future studies could test these predictions by investigating the genetic architecture of loci involved in TGP and selection on expression of parentally transmitted information in both sexes. The diversity of listening outcomes generated by our model is consistent with the diverse patterns of TGP seen in nature across a wide range of taxa (Mousseau and Fox 1998; Agrawal et al. 1999; Galloway 2001; Anway et al. 2005; Pembrey et al. 2006; Galloway and Etterson 2007; Dunn et al. 2011; Salinas and Munch 2012; Ducatez et al. 2012; Babenko et al. 2015; Akkerman et al. 2016; Guillaume et al. 2016; Emborski and Mikheyev 2019). Our model therefore provides a unifying framework for understanding the origin and maintenance of diversity in observed patterns of TGP.

## DATA ACCESSIBILITY

The simulation data that support the findings of this study are available on Dryad Data Repository: https://datadryad.org/stash/share/a_c-2-t-hugOmUf0IdBKmw6gcL8ujoC_eDKVovj39fA The IBM code that generated the data is also available via this link.

---

**SUPPLEMENTARY FIGURE S1.**
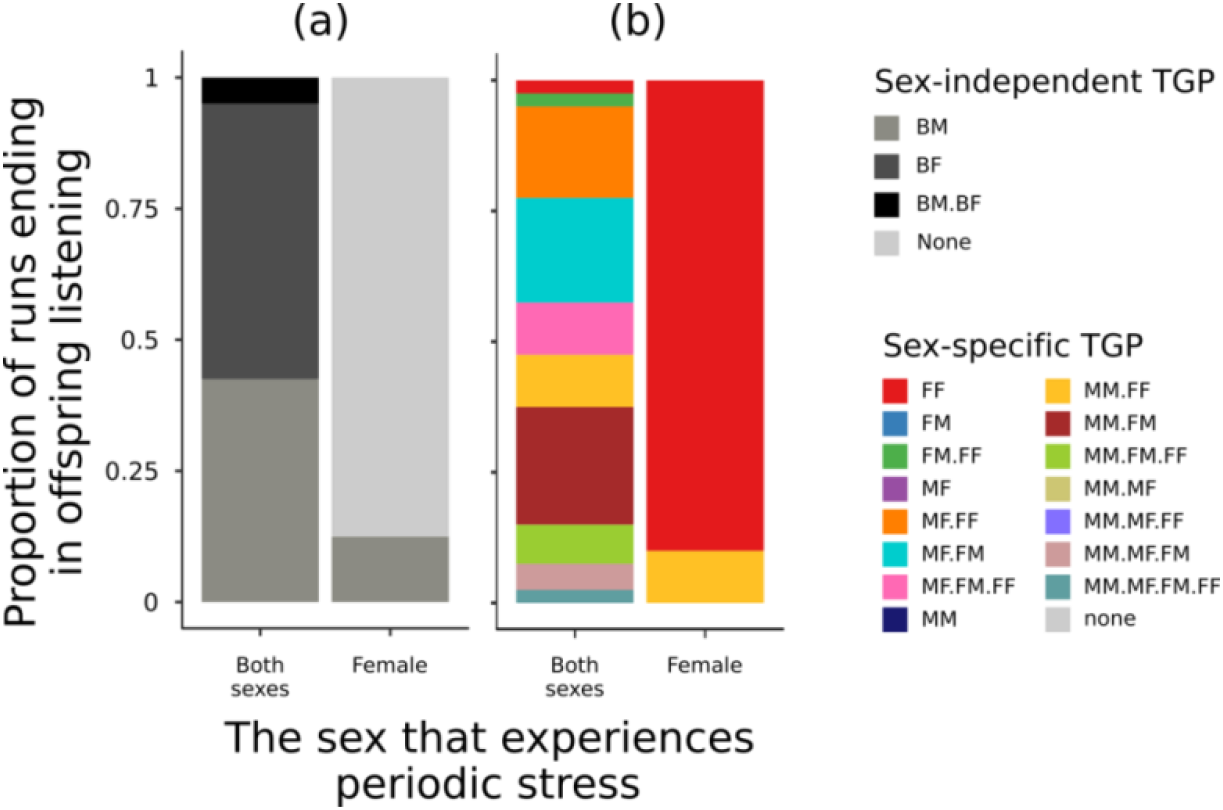
Sex-independent listening (a) and sex-specific listening (b) when environmental stress is periodic. Fluctuating stress generates patterns of TGP that are very similar to those resulting from a single, extended bout of elevated stress (see main text). The plot shows the proportion of 40 simulation runs ending in listening outcomes at generation 1000 when conditions alternate between benign and stressful every 25 generations, causing females or both sexes to experience fluctuating stress. Other settings are: ‘best male’ version of each model; mating neighbourhood size = 25 patches.

**SUPPLEMENTARY FIGURE S2.**
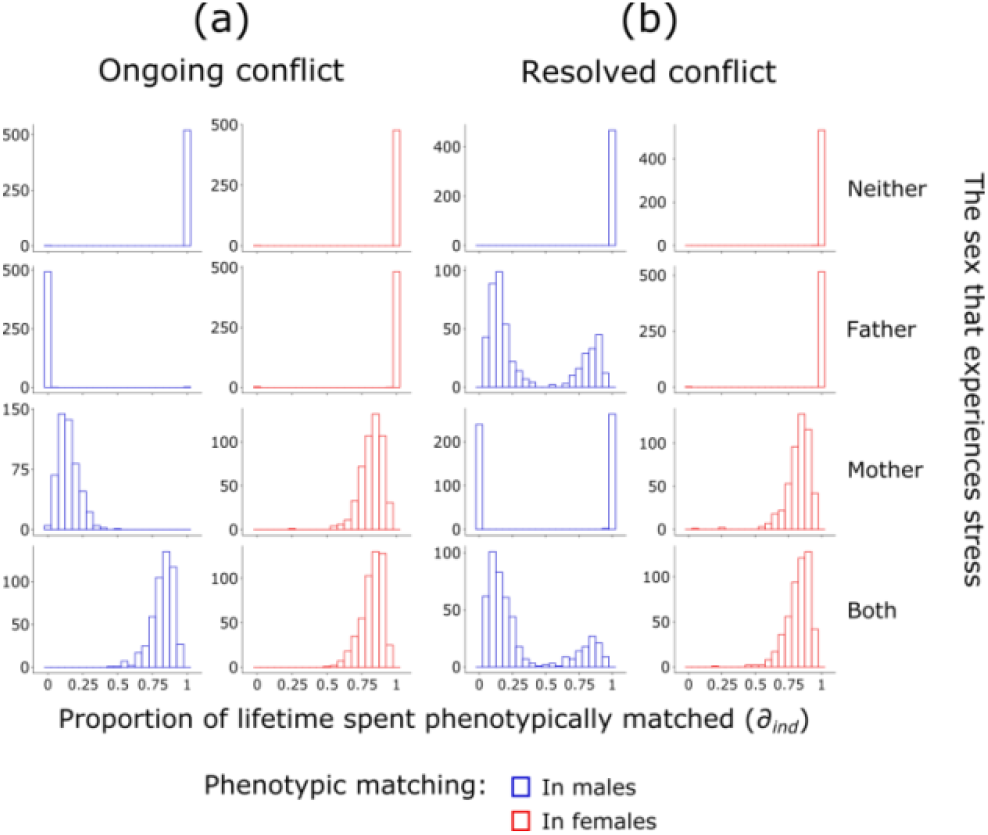
Patterns of phenotypic matching (*∂*_*ind*_) when mating is random. Histograms show the outcome of selection on males (blue) and females (red) at the end of a single simulation run of the ‘random mating’ version of the ongoing conflict model (a) and resolved conflict model (b) at generation 1000. The absence of sexual selection on males allows fecundity selection to maximise female matching in (a) regardless of which parent experiences stress, and limits maximal matching in males in (b). For the unstressed sex in both (a) and (b), distributions of *∂*_*ind*_ show little variation around the peak because individuals of this sex rarely encounter more conspecifics than their sex’s encounter threshold (*α*_*m*_ or *α*_*f*_), and so are almost always uniformly well-matched. Other settings: mating neighbourhood size = 25 patches.

**SUPPLEMENTARY FIGURE S3.**
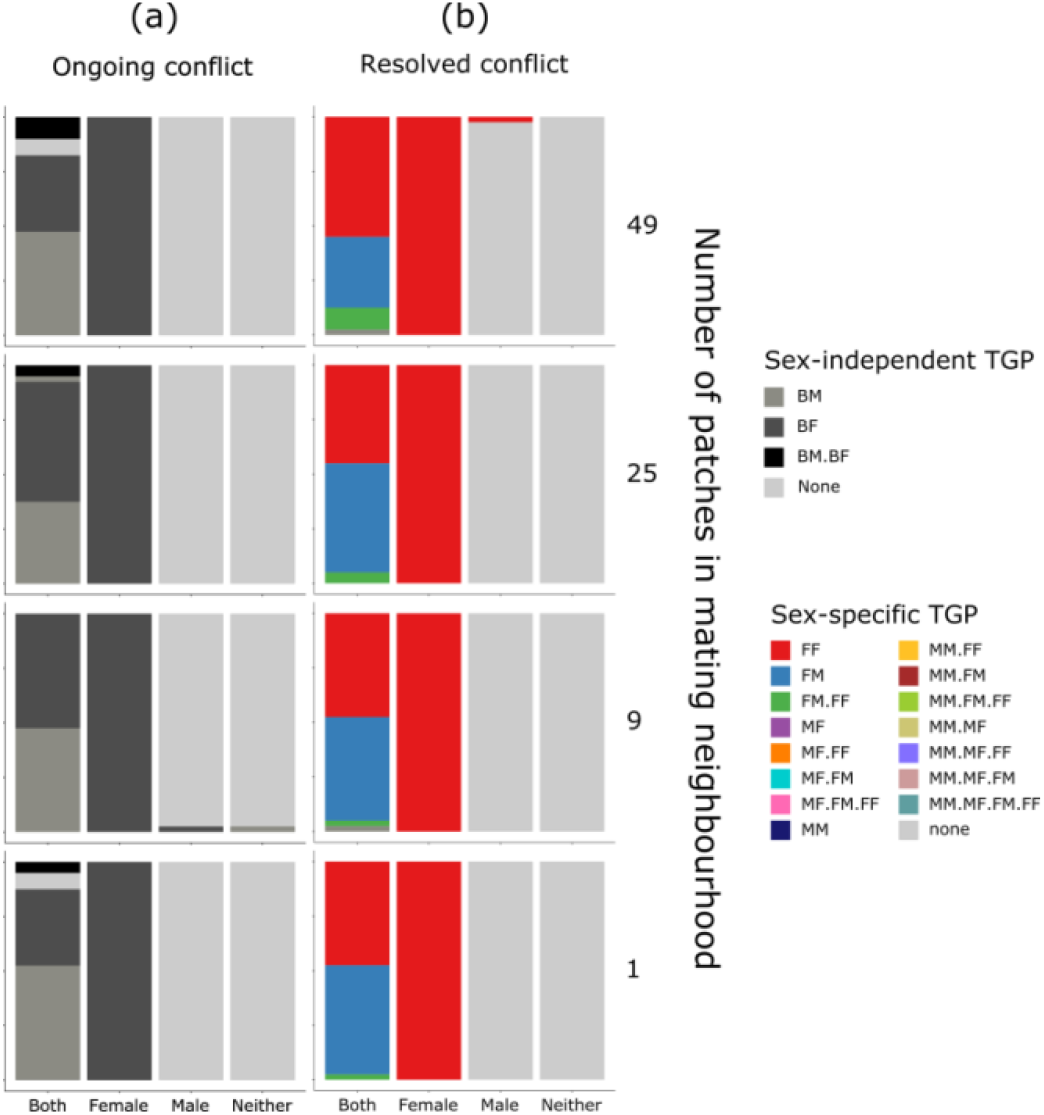
Sex-independent listening (a) and sex-specific listening (b) when mating is random. Graphs on the left (a) show the proportion of runs of the random mating version of the ongoing conflict model ending in sex-independent paternal TGP (dark grey), maternal TGP (medium grey), both types of TGP (black), and no TGP (light grey) at generation 1000. Graphs on the right (b) show the proportion of simulation runs of the random mating version of the resolved conflict model ending in sex-specific TGP at generation 1000. Colours in the legend represent all possible combinations of listening. The first letter in each double-letter code represents the sex of the offspring that does the listening (M = males (i.e., sons); F = females (i.e., daughters); B = both sexes), and the second letter represents the sex of the parent that is listened to (M = males (i.e., fathers); F = females (i.e., mothers)). Thus, for example, BM indicates a sex-independent paternal effect (i.e., sons and daughters both listening to fathers), and MF indicates a son-specific maternal effect (i.e., sons listening to mothers). Colours with more than one double-letter code indicate runs where more than one combination of sex-independent or sex-specific TGP evolves. Runs were counted as ending in TGP if listening allele frequencies were ≥ 0.95.

